# Capturing the dynamics of peripersonal space by integrating sound propagation properties and expectancy effects

**DOI:** 10.1101/756783

**Authors:** Lise Hobeika, Marine Taffou, Thibaut Carpentier, Olivier Warusfel, Isabelle Viaud-Delmon

## Abstract

**Highlights:**

- Logarithmically distributed auditory distances provides an apt granularity of PPS
- Measuring expectation helps to interpret behavioral impact of audiotactile integration
- Tactile RTs follows a logarithmic decrease due to audiotactile integration
- Peripersonal space is better characterized and quantified with this refinement

**Background:** Humans perceive near space and far space differently. Peripersonal space, i.e. the space directly surrounding the body, is often studied using paradigms based on auditory-tactile integration. In these paradigms, reaction time to a tactile stimulus is measured in the presence of a concurrent auditory looming stimulus.

**New Method:** We propose here to refine the experimental procedure considering sound propagation properties in order to improve granularity and relevance of auditory-tactile integration measures. We used a logarithmic distribution of distances for this purpose. We also want to disentangle behavioral contributions of the targeted audiotactile integration mechanisms from expectancy effects. To this aim, we added to the protocol a baseline with a fixed sound distance.

**Results:** Expectation contributed significantly to overall behavioral responses. Subtracting it isolated the audiotactile effect due to the stimulus proximity. This revealed that audiotactile integration effects have to be tested on a logarithmic scale of distances, and that they follow a linear variation on this scale.

**Comparison with Existing Method(s):** The granularity of the current method is more relevant, providing higher spatial resolution in the vicinity of the body. Furthermore, most of the existing methods propose a sigmoid fitting, which rests on the intuitive framework that PPS is an in-or-out zone. Our results suggest that behavioral effects follow a logarithmic decrease, thus a response graduated in space.

**Conclusions:** The proposed protocol design and method of analysis contribute to refine the experimental investigation of the factors influencing and modifying multisensory integration phenomena in the space surrounding the body.

## 1. Introduction

The space around the body, called peripersonal space (PPS), is selectively encoded in the brain by a neural network linked to multisensory integration processes (Bernasconi et al., 2018; Colby et al., 1993; Farnè and Làdavas, 2002; Gentilucci et al., 1988; Graziano and Cooke, 2006; Graziano and Gross, 1993; Làdavas and Farnè, 2004). Whereas PPS multisensory encoding has been examined in monkeys and patients with neurophysiological invasive methods, developing a behavioral method to study PPS in healthy humans is not straightforward (Cléry and Ben Hamed, 2018). Following a large literature describing behavioral effects of multisensory integration as a function of the spatial proximity of sensory stimuli to the body (Ladavas et al., 2001; Maravita et al., 2003), Canzoneri and colleagues’ developed an audiotactile method to study PPS in humans (Canzoneri et al., 2012). In this method, participants perform a speeded tactile detection task while an irrelevant sound is looming towards them from the frontal hemifield. The tactile stimulus is delivered at the beginning of the sound, i.e. when the sound source is far from the participant body, or at the end of the sound, i.e. when the sound source is near the participant body, or at intermediate positions in space. The analysis of participants’ tactile detection time as a function of the sound source position in space gives information on PPS morphometry.

This method has several assets. First, it is an implicit measure: participants are not aware that the sound position can modulate their reaction time. Second, the method rests on multisensory reaction time facilitation effects with looming stimuli. Thus, the protocol is based on multisensory integration behavioral effects that have been extensively studied (Hershenson, 1962; Spence et al., 1998; Sumby and Pollack, 1954) and on moving looming stimuli, which are particularly relevant for the study of PPS. PPS multisensory neurons in monkeys are indeed very reactive to movements, and especially to movements looming towards the body (Colby et al., 1993; Graziano and Gross, 1993; Rizzolatti et al., 1981). The relevance of looming stimuli has also been evidenced in humans (Cléry et al., 2015; Kandula et al., 2015; Tajadura-Jiménez et al., 2010; Van der Biest et al., 2016). Lastly, this method measures multisensory facilitation at multiple positions in space, which gives a higher spatial resolution of PPS morphometry. However, the method has some limitations linked to the involvement of auditory dynamic cues and expectancy effects.

Localizing a sound in 3D space is a complex cognitive process that takes into account a wide variety of acoustic parameters. Two of the main acoustic cues for the perception of distances are sound level change and reverberation (Shinn-Cunningham, 2000; Zahorik et al., 2005). Sound intensity globally decreases when sound source distance increases. In open-air condition, the sound intensity follows an inverse-square law, which implies an intensity decrease of 6 dB per doubling of source distance. The sound intensity logarithm (in dB) varies linearly with the logarithm of distances. The sound level is a relative cue for distance perception: it does not allow an estimation of the absolute distance to the source, as a modulation in sound level could be due to a moving sound source or a to decrease of the intensity of the emitted sound. Distance perception is also based on the ratio between the levels of the reverberated and of the direct sound, called direct-to-reverberant sound energy ratio. Sound reverberation is caused by the multiple reflections of the acoustic waves occurring on obstacles and space boundaries, and creates a diffuse surrounding sound field. In contrast with the level of the direct sound, in a closed space, the level of the reverberated sound field does not depend on the distance to the source. Consequently, the direct-to-reverberant sound energy gives an absolute cue on the sound-source distance. Combining those two cues, participants are usually good at comparing the position of two sound sources in space, but are not highly accurate in the evaluation of absolute distances (Kolarik et al., 2015; Zahorik et al., 2005).

Overall, distance perception is mainly based on direct sound level, which varies logarithmically with sound distances. Studies on source distance perception usually test distances distributed regularly on a logarithmic scale (e.g. Alais and Carlile, 2005; Fontana and Rocchesso, 2008; Zahorik and Wightman, 2001). However, to our knowledge, PPS studies systematically test distances distributed linearly. With this method, the perceptual differences between distances are highly irregular. For example, there are small acoustic differences when a sound source moves from 6m20 to 5m20 from the observer, but there is a larger one for a movement from 1m20 to 0m20. Thus, using distances spaced regularly on a logarithmic scale should be more relevant to study the extent of PPS, as it equalizes the perceptual differences between each tested distance. A logarithmic based distances distribution should provide a better granularity of the dynamics of multisensory integration processes in space.

Another issue relies on expectation mechanisms. In Canzoneri’s paradigm, a tactile stimulus is presented during a three second sound in the majority of the experimental trials, and there is no more than one tactile stimulus per sound. Thus, there is a high probability that a tactile event occurs during the duration of the sound. If no tactile event arrived yet after a two-second duration of sound, the probability that a tactile event occurs in the last second of the sound is even higher. Therefore, the probability of the occurrence of a tactile event evolves during the duration of the sound with the delay from the sound onset. As the distance of the sound from the body also evolves with the delay from sound onset, the temporal expectancy effect is entangled with the changes in sound distance. Consequently, it is not possible to understand the respective role of expectation and sound distance in the behavioral effects observed in the presence of the looming sounds. A study by Kandula and colleagues focused on the contribution of tactile expectancy on RTs in Canzoneri’s method (Kandula et al., 2017). They demonstrated that when the probability to receive a tactile stimulation at each trial is high, the large change in expectancy during a trial impacts RTs and masks the multisensory integration effects. It is important to note that expectation is not entirely responsible for the results on PPS obtained with this protocol. If RTs variations were explained only by expectancy effects, the same results should be found for looming and receding sounds which is not the case (Canzoneri et al., 2012; Serino et al., 2015).

A solution to help disentangling the contribution of expectation from audiotactile integration effects on RTs is to add a baseline, in which there is expectancy but little audiotactile integration. Measuring tactile RTs using a sound fixed in space and located far from participants’ body gives a temporal reference for participants to expect a tactile stimulation. In this case, the observed behavioral effects cannot be related to a modulation of sound distance, hence not to multisensory integration.

Lastly, we want to obtain an efficient description of the evolution of audiotactile integration in space. Some studies used a sigmoidal fitting to describe RTs evolution with sound source distance in space (e.g. Canzoneri et al., 2012; Ferri et al., 2015; Taffou and Viaud-Delmon, 2014), some used a linear fitting (e.g. Cardini et al., 2019; Salomon et al., 2017) and others did not performed data fitting (e.g. Hobeika et al., 2019; Noel et al., 2015b; Serino et al., 2015). As there is no clear consensus on the most suited manner to describe and fit this kind of data, it would be interesting to examine audiotactile integration effects on RTs as a function of the sound source distance when auditory distance and expectation mechanisms are taken into account.

The aim of the present study is to provide a better description of peripersonal space morphometry, using Canzoneri’s and colleagues audiotactile task (Canzoneri et al., 2012). First, to measure the impact of expectancy effect on RTs during a trial, we added a condition, in which the sound source is fixed in space and located at the starting distance of the sound source of the looming sound. Tactile stimulations occurred at the same delays from sound onset in both the fixed sound and the looming sound conditions. Second, we modified the distribution of tested distances in space. As auditory perception in depth strongly involves loudness, which rests on a logarithmic process, it should be more relevant to test audiotactile integration at distances spaced regularly on a logarithmic scale. To test this hypothesis, participants completed two sessions of test: one with distances spaced regularly on a linear scale, and one with distances spaced regularly on a logarithmic scale. Finally, we tried to find a simple fitting representing the evolution of RTs with sound distances. Additionally, we introduced a test at the end of the experiment, in which we assessed participants’ emotional response to the looming sound according to its position in depth. This test was based on the Behavioral Assessment Test (BAT), which is widely used in clinical psychology to assess the level of fear of patients in response to a phobic object that is getting closer and closer to them (e.g. Lang and Lazovik, 1963; Van Bockstaele et al., 2011). In the present study, the test was an auditory BAT (aBAT) that provided an indirect assessment of the perceived position of the sound, and examined how the emotional value attributed to the sound distance evolves according to a logarithmic and linear repartition of distances

## 2. Material and methods

### 2.1 Participants

Nineteen healthy individuals (12 females, age 24.7 ± 3.6) with normal audition and touch took part in the study. Sample size was decided a priori based on previous studies using similar audiotactile paradigms (Serino et al., 2015; Taffou and Viaud-Delmon, 2014). All of them were right-handed. As PPS is linked to handedness (Hobeika et al., 2018), participants’ handedness was verified with a questionnaire measuring skilled hand preference. The scores on this questionnaire, called the Flinders Handedness survey (FLANDERS) (Nicholls et al., 2013), range from -10 for strong left-handed individuals to +10 for strong right-handed individuals. Participants received a financial compensation of 10€/hour for their participation. They provided a written informed consent prior to the experiment, which was approved by the Institutional Review Board of the French National Institute of Health and Medical Research (INSERM, IRB00003888).

### 2.2 Apparatus

Participants sat on a chair in a soundproofed room. Both of their hands were palms-down on a table, in contact with their body and aligned with their mid-sagittal plane. A black fabric hid participants’ hands. Participants were equipped with Beyer Dynamic DT770 headphones. To control for the visual stimulation and gaze direction, participants were instructed to fix a permanent visual cross located at 1.5m in front of them at eye level.

The auditory stimuli were sounds composed of bursts train (44100 Hz digitization). The bursts train consisted of a succession of Gaussian white noise bursts equalized in intensity (created with Matlab®). Each burst lasted 30ms with 10ms rise and fall times. The time interval between two bursts was 65ms. For the purpose of the experiment, two types of spatialized auditory stimulus were created based on this bursts train sound: static auditory stimuli (fixed sounds) and dynamic auditory stimuli (looming sounds). In order to simulate spatialized auditory sources, the bursts train sound was processed through binaural rendering in the Max/MSP (6.1.10) environment using the Spat library (Carpentier et al., 2015). Extra time was left after the last burst (115ms) in order to account for the reverb tail. The simulation consisted in a sound source placed in a virtual shoebox room (685m^3^), which first reflections up to order 3 and late reverberation were rendered dynamically. The spatialization of the direct sound and of the first reflections were rendered using non-individual head related transfer functions (HRTF) taken from the LISTEN HRTF database (http://recherche.ircam.fr/equipes/salles/listen/). With this procedure, the virtual sound source location can be manipulated by rendering accurate auditory cues such as frequency spectrum, intensity, and inter-aural differences. For both types of auditory stimuli, we simulated a sound source located in the frontal hemifield in the right hemispace (azimuth -60°). All participants confirmed that they could clearly locate sound sources in the right hemispace. For the fixed sounds, the virtual sound source was static and located at 640cm from participants’ head center, at ear level. For the looming sound, the virtual sound source was approaching participants’ head center from 640cm to 20cm, at ear level and at a constant speed (2.1 m.s^-1^).

The tactile stimulus was a vibratory stimulus delivered by means of a 28mm miniature loudspeaker on the palmar surface of the left index finger of participants. A sinusoid signal was emitted for 20ms at 250 Hz. With these parameters, the vibration of the loudspeaker was perceivable, but the sound was inaudible. A PC running Presentation® software was used to control the presentation of the stimuli and to record the responses.

### 2.3 Design and procedure

The experiment was composed of two experimental sessions. Participants completed the two sessions on two different days. The first session consisted of an audiotactile test. The second session consisted of an audiotactile test followed by three auditory Behavioral Assessment Tests (aBATs). The aim of the audiotactile tests was to evaluate the advantages of our protocol modifications, in terms of expectancy assessment and granularity of measures, for the study of audiotactile integration behavioral consequences with sound distances. The aim of the aBATs was to indirectly assess the perceived position of the sound sources by examining the emotional value attributed to the sound according to its distance from participants.

#### 2.3.1 Audiotactile tests

At the beginning of the experimental session, participants were asked to place their left index finger on the vibrator and instructed to press a button with their right finger each time they detected a tactile stimulus. For each trial, an auditory stimulus (either a fixed or a looming sound) was presented for 3250ms. The auditory stimulus was preceded by 300ms of silence. A period of silence, with a duration randomly varying between 700 and 1100ms, also occurred after the offset of the sound. In 66.6% of the trials (experimental trials), a tactile stimulus was presented along with the auditory stimuli. The remaining 33.3% trials were catch trials with auditory stimulation only. Participants were instructed to ignore the auditory stimuli and to respond as quickly as possible to the tactile stimulation. They were asked to emphasize speed, but to refrain from anticipating. Reaction times (RTs) were measured.

During experimental trials, tactile stimulations were delivered at different delays starting from sound onset. We presented the tactile stimulation 10ms after the burst onset whichever burst among the 33 bursts of the sound was targeted in order to apply the desired delay. When the auditory stimulus presented during the trial was a fixed sound, the distance of the virtual sound source was 640cm when tactile stimulation occurred, regardless of the delay applied between sound onset and tactile stimulation. In contrast, when the auditory stimulus presented during the trial was a looming sound, the distance of the virtual sound source when tactile stimulation occurred depended of the delay applied between sound onset and tactile stimulation – far distances for low temporal delays and near distances for high temporal delays – (see **Figure 1**).

**Figure 1:**
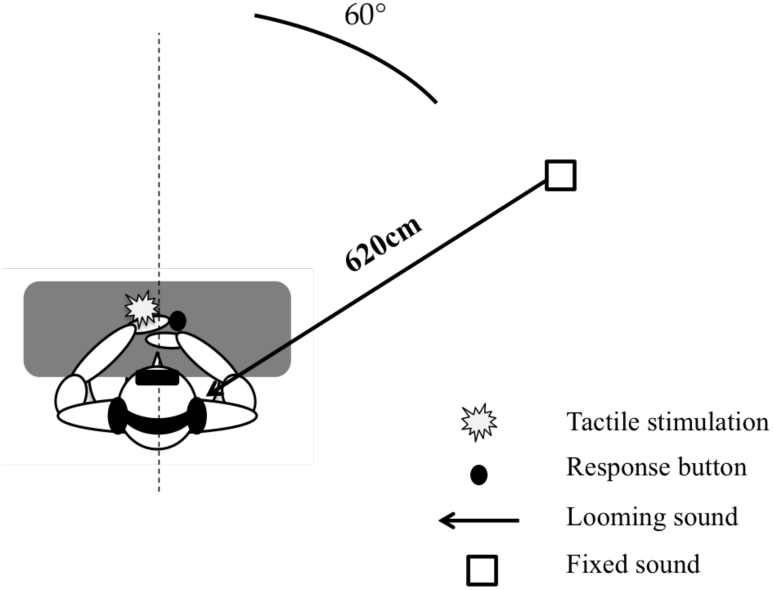
Participants performed the audiotactile task by responding to a tactile stimulation on their hand while a task-irrelevant spatialized sound was presented to them in their right hemispace. The sound could be moving towards them from 640 cm to 20cm distance (looming sound), or be static and located at 640cm from the center of the participants’ head (fixed sound)

Two sets of six distances with different spatial distributions in depth were tested in two separated experimental sessions: a set with a logarithmic distribution of distances (640cm, 320cm, 160cm, 80cm, 40cm, 20cm) and a set with a linear distribution of distances^1^ (640cm, 520cm, 400cm, 260cm, 140cm, 20cm). The temporal delays used for tactile stimulus delivery and the corresponding distances of the looming sound source at delivery are described in **Table 1** for both distances distributions. The evolution of sound level with the approach of the sound source is described on **Figure 2, panels a and d**. Values are given with reference to the direct sound level at the closest distance, i.e. 20cm. According to the direction of incidence, the sound pressure level delivered to the ears of the participant ranged from 42 to 69dBA and from 45 to 73dBA, for the contralateral and the ipsilateral ears respectively. As depicted by the **panels b and e of Figure 2**, whereas the difference of sound level between distance conditions are identical when the distances tested are distributed logarithmically, this is not the case when the distance conditions are distributed linearly.

**Table 1.** Description of the delays of tactile stimulation from sound onset used in each distance distribution sessions, and the corresponding distances for bimodal trials, for the looming sound and the fixed sound.

| | $T_{before}$ | T1 | T2 | T3 | T4 | T5 | T6 | $T_{after}$ |
| --- | --- | --- | --- | --- | --- | --- | --- | --- |
| **Logarithmic distribution** |  |  |  |  |  |  |  |  |
| Delay from sound onset (ms) | 50 | 405 | 1925 | 2685 | 3065 | 3255 | 3350 | 3800 |
| Looming sound distance (cm) |  | 640 | 320 | 160 | 80 | 40 | 20 |  |
| Fixed sound distance (cm) |  | 640 | 640 | 640 | 640 | 640 | 640 |  |
| **Linear distribution** |  |  |  |  |  |  |  |  |
| Delay from sound onset (ms) | 50 | 405 | 975 | 1545 | 2210 | 2780 | 3350 | 3800 |
| Looming sound distance (cm) |  | 640 | 520 | 400 | 260 | 140 | 20 |  |
| Fixed sound distance (cm) |  | 640 | 640 | 640 | 640 | 640 | 640 |  |

**Figure 2.**
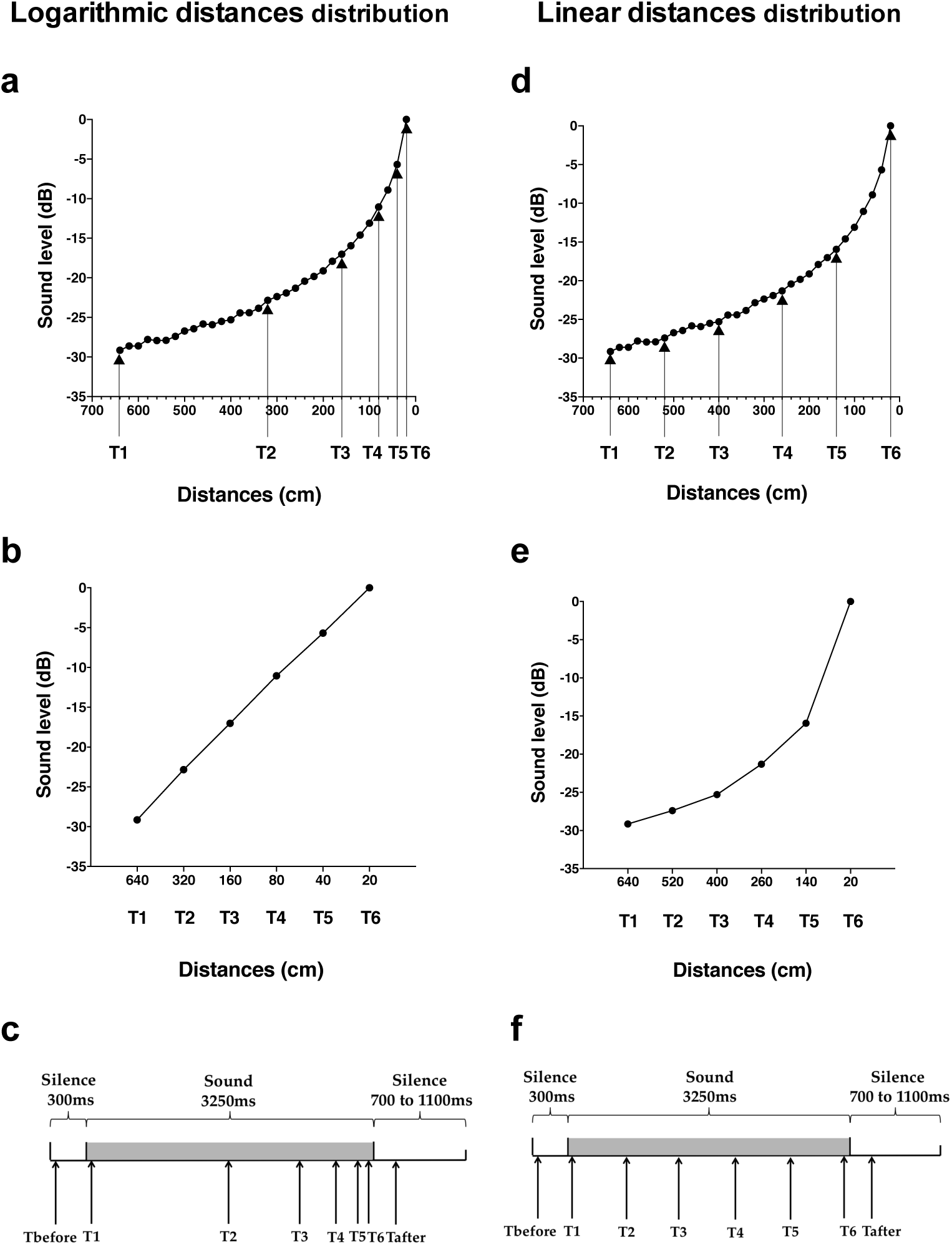
**(a, b, d, e)** Sound level at each burst of the looming sound as a function of the tested distances. Values are given with reference to the direct sound level at the closest distance, i.e. 20cm. Arrows indicate tested distances in the logarithmic distances distribution **(a)** and in the linear distances distribution (**d)**. As illustrated on graphs **a** and **d**, sound level varies exponentially with distance. As expected, sound level varies linearly when distances are regularly spaced on a logarithmic scale (see **b**), giving a relevant mapping of space. When distances are spaced regularly on a linear scale (see **e**), the sound level variation is exponential. In this linear distribution, the measures at far distances may be redundant, whereas there is a lack of precision for close distances**. (c, f)** Description of a trial. For each trial, one tactile stimulation was delivered at one among eight possible delays from sound onset (Tbefore, T1 to T6 and Tafter), corresponding to six possible distances of the sound source from participants’ body and to two unimodal trials in the logarithmic distances distribution **(c)**, and in the linear distances distribution **(f)**

In order to measure RTs in the unimodal tactile condition (without any sound), tactile stimulations were also delivered during the silent periods, preceding or following sound administration, namely at –250ms (Tbefore) and at 3800ms (Tafter) from sound onset (see a temporal description of trials in the logarithmic experimental session and in the linear experimental session in **Figure 2c and 2f** respectively).

Each participant completed both experimental sessions: one with a logarithmic distances distribution and one with a linear distances distribution. The order of the sessions was counterbalanced between participants. Both sessions were realized in two different days, separated by a maximum of three days.

The total test consisted of a random combination of 24 repetitions of the target stimuli in each of the 32 conditions. The within factors were: DISTANCES DISTRIBUTIONS (two levels: logarithmic/linear), SOUND MOUVEMENT (two levels: looming/fixed) and DELAY (eight levels: Tbefore, T1, T2, T3, T4, T5, T6 and Tafter). In both experimental sessions, trials were equally divided in 8 blocks of 72 trials, lasting about 5 min each. Each block contained 48 trials with a tactile target, randomly intermingled with 24 catch trials.

#### 2.3.2 Auditory Behavioral Assessment Tests

Participants were seated on a chair and equipped with Beyer Dynamic DT770 headphones. They were instructed to look at the fixation cross located at 1m50 in front of them, at eye level. During each aBAT, a sound was played continuously through participants’ headphones. Participants performed three different aBATs: the Logarithmic aBAT, the Linear aBAT and the fixed aBAT. Each aBAT consisted of six steps. The sound distance at each step corresponded to the sound distance of the corresponding audiotactile tests. In the logarithmic aBAT, sound source distances at each step were spaced regularly on a logarithmic scale (640cm, 320cm, 160cm, 80cm, 40cm, 20cm); in the Linear aBAT, sound source distances at each step were spaced regularly on a linear scale (640cm, 520cm, 400cm, 260cm, 140cm, 20cm); and in the Fixed aBAT, the sound source was static, located at 640cm at each step. The emotional experience induced by the sound as a function of its distance was assessed with Subjective Units of Distress (SUD; Wolpe, 1973). SUD is a self-report typically used for measurement of experienced fear or discomfort, which has been shown to correlate with physiological measures of arousal state (Thyer et al., 1984). At each step of the aBATs, participants had to rate their level of discomfort with SUD on a scale from 0 to 10 - 0 corresponding to an absence of discomfort and 10 to the worst discomfort possible -. Between each step, the sound source moved from the former position to the next step position at a regular speed for 2 seconds. 500ms after the sound source reached its targeted position, a sound signal was played to indicate participants that they had to rate their level of discomfort. The order of presentation of the Logarithmic aBAT and Linear aBAT was counterbalanced between participants; the fixed aBAT was always presented at the end.

## 3. Results

One participant was excluded from analysis due to a high rate of misses in the audiotactile test (23.8% of miss, m±sd of the sample: 2.6 ± 2.0 % of miss). There were eighteen remaining subjects (11 females, age 25.0 ± 3.5). All remaining participants were right-handed, as verified by their FLANDERS score (range: from 6 to 10; *M* ± *SD* = 9.6 ± 1.1).

### 3.1 Audiotactile tests

The performances were analyzed in terms of RTs. We considered as a valid answer all RTs between 100 and 1000ms after stimulus onset. For the analyses, RTs were averaged for each subject and for each of the 32 conditions separately (2 DISTANCES DISTRIBUTION * 2 SOUND MOVEMENT * 8 DELAYS).

We conducted an ANOVA on the mean RTs, with the within-subject factors DISTANCES DISTRIBUTION (two levels: logarithmic, linear), SOUND MOVEMENT (two levels: fixed, looming), and the factor DELAY (eight levels: Tbefore, T1, T2, T3, T4, T5, T6, and Tafter). Analysis revealed a significant main effect of SOUND MOVEMENT (F_(1,17)_ = 46.7, p < .001, *η _p_* ^*2*^ = .733), of DELAY (F_(7,119)_ = 48.8, p < .001, *η*_*p*_^*2*^ = .720), and significant two-way interactions between the factors DISTANCES DISTRIBUTION x MOVEMENT (F_(1,17)_ = 36.4, p < .001, *η*_*p*_^*2*^ = .681), DISTANCES DISTRIBUTION x DELAY (F_(7,119)_ = 11.1, p < .001, *η*_*p*_^*2*^ = .394) and MOVEMENT x DELAY (F_(7,119)_ = 12.2, p < .001, *η*_*p*_^*2*^ = .419). Finally, the three-way interaction DISTANCES DISTRIBUTION × MOVEMENT × DELAY was significant (F_(7,119)_ = 3.34, p < .01, *η*_*p*_^*2*^ = .164), suggesting that RTs were impacted differently by the sound movement depending on the distances distribution.

#### 3.1.1 Assessment of the expectancy with the fixed sound condition

We tested whether the RTs in the fixed sound condition can be used as a baseline to measure the tactile expectancy effects. To this aim, we first compared RTs on unimodal trials to bimodal ones in the fixed sound condition, in both distances distributions. We then analyzed if the movement of the sound impacted unimodal tactile detection when it was presented after the sound. We performed those analysis using post-hoc tests in accordance with the significant three-way interaction DISTANCES DISTRIBUTION x MOVEMENT x DELAY described in the previous paragraph. Finally, we fitted the bimodal data of the fixed sound condition as a function of the delay to have a description of the evolution of expectancy effects.

**Comparison of unimodal trials and bimodal trials for fixed sound trials.** In both distances distribution conditions, RTs occurring at Tbefore were significantly slower than RTs occurring at T2, T3, T4, T5, T6 and Tafter (logarithmic distances distribution: Post-hoc Newman-Keuls’ test: p <.001 in all cases; linear distances distribution: Post-hoc Newman-Keuls’ test: p <.05 in all cases). RTs occurring at Tafter delay were significantly faster than RTs at Tbefore, T1 and T2 (logarithmic distances distribution: Post-hoc Newman-Keuls’ test: p <.001 in all cases; linear distances distribution: Post-hoc Newman-Keuls’ test: p <.05 in all cases) (see **Figure 3**). Those results indicate that there was a decrease of RTs with the factor delay, likely due to expectancy effects. There was no significant difference between Tbefore and T1, and between T6 and Tafter, suggesting that there was no audiotactile integration with the fixed sound, located far from the body (640cm).

**Figure 3.**
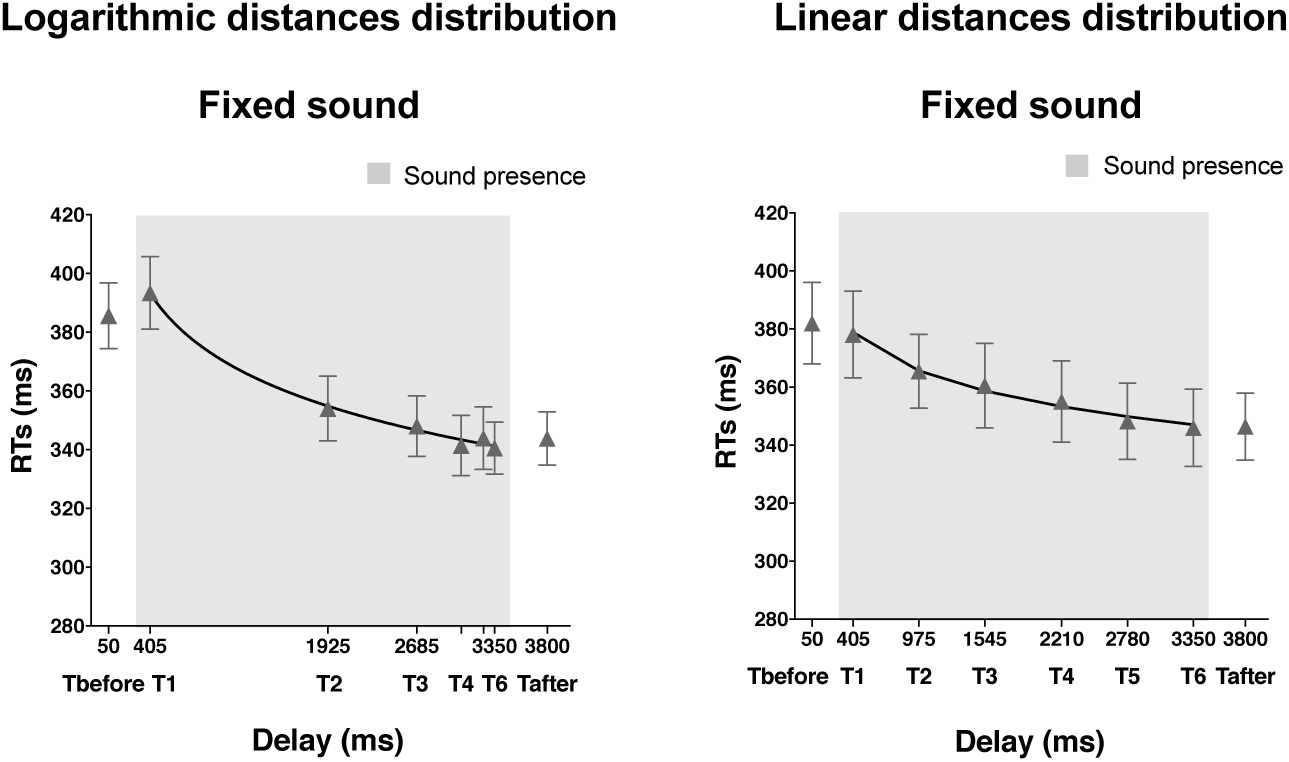
Analysis of the expectancy effect. This figure reports the mean tactile reaction times (±SEM) as a function of the delay of tactile stimulation delivery from the trial beginning, in presence of a sound fixed in space (located at 640cm from participants’ head center). Tactile stimulation occurred alone in the unimodal trials (Tbefore, Tafter), or occurred in presence of a sound in the bimodal trials (T1, T2, T3, T4, T5 and T6). The shaded region indicated the duration of the sound. RTs are fitted with a decreasing logarithmic function on both figures.

**Effect of sound movement on unimodal trials after the sound:** for both distances distributions, sound movement did not significantly impact RTs at Tafter (logarithmic distances distribution: Post-hoc Newman-Keuls’ test: p= 0.4, Tafter_fixed_sound: 343.8 ± 9.1 ms, Tafter_looming_sound: 335.7 ± 6.7 ms ; linear distances distribution: Post-hoc Newman-Keuls’ test: p= 0.3, Tafter_fixed_sound: 346.4 ± 11.6 ms, Tafter_looming_sound: 335.1 ± 10.1 ms). Those results suggest that the presence of a fixed or a looming sound had no post-effect on the detection of the unimodal tactile stimulation at Tafter.

**Description of the expectancy effects.** In order to describe the effects of expectancy on tactile RTs, we fitted RTs from the bimodal trials in the fixed sound condition with two different functions. We hypothesized that the evolution of tactile RTs as a function of the delay of tactile delivery could follow a linear decrease or a logarithmic decrease, depending on the distances distribution. To test this hypothesis, we plotted participants’ RTs in the bimodal trials in the fixed sound condition as a function of the delay of tactile delivery, in the logarithmic and linear distances distribution sessions. Then, we fitted a linear and a logarithmic function to the data, separately for each participant. We used, as the logarithmic function *y(x) = a.log(x) + b*, where *x* represented the independent variable (i.e. the delay of tactile stimulation from sound onset), *y* the dependent variable (i.e. the RTs), *a* theslopeand*b*thethresholdfor*x=1*.Thelinear function we used, is described by *y(x) = a.x + b* where *a* is the slope and *b* is the intercept at *x = 0*. As both functions contain two parameters, we could compare the quality of the fitting by directly comparing the root mean square errors values (RMSE) for each participant. For the logarithmic distances distribution, analysis revealed that the logarithmic function was significantly better to describe participants’ RTs than the linear function (t(17) = -2.27, p <.05, two-tailed, RMSE_logarithmic_fitting_= 9.31, RMSE_Linear_fitting_ = 10.48) (see **Figure 3**). For the linear distances distribution, both functions gave the same performance at describing participants’ RTs (p >.05, two-tailed, RMSE_logarithmic_fiting_ = 11.1, RMSE_Linear_fitting_ = 11.2).

We then compared the intensity of the expectancy effects for both distances distributions. To this aim, we compared the parameters obtained with the logarithmic decrease fitting for both distances distributions. We found that both parameters **a** and **b** were impacted by the distances distribution (**parameter a:** t(17) = -2.44, p <.05, two-tailed, aLogarithmic_distribution= -24.6, aLinear_distribution = -15.1; **parameter b:** t(17) = -2.10, p = .05, two-tailed, bLogarithmic_distribution= 541.0, bLinear_distribution = 496.5). Parameters *a* and *b* of the logarithmic function were significantly larger in the logarithmic distances distribution than in the linear one. These findings indicate that the evolution of the expectancy effect with time depended on the tested delays of tactile stimulation from sound onset.

#### 3.1.2. Evaluation of the audiotactile integration effect on RTs in the linear and logarithmic distances distributions

We analyzed the effects of distances distribution, sound movement and delay on RTs. Considering that the three-way interaction DISTANCES DISTRIBUTION × SOUND MOVEMENT × DELAY was significant (see above), we performed separated ANOVAs to analyze the interaction between sound movement and delay for each distances distribution independently, on bimodal trials only.

**Logarithmic distances distribution.** We conducted an ANOVA on the mean RTs of the Logarithmic distances distribution on bimodal trials only, with the within-subject factors SOUND MOVEMENT (two levels: fixed, looming), and the factor DELAY (six levels: T1, T2, T3, T4, T5 and T6). Analysis revealed a significant main effect of SOUND MOVEMENT (F_(1,17)_ = 76.3, p < .001, *η*_*p*_^*2*^ = .818),and a significant main effect of DELAY (F_(5,85)_ = 49.2, p < .001, *η*_*p*_^*2*^ = .743). The analysis also revealed that the two-way interaction SOUND MOVEMENT x DELAY was significant (F_(5,85)_ = 18.2, p < .001, *η*_*p*_^*2*^ = .518), indicating that tactile RTs variation with the temporal delay of tactile delivery from sound onset depended on the movement of the sound (see **Figure 4 left figure**).

**Figure 4.**
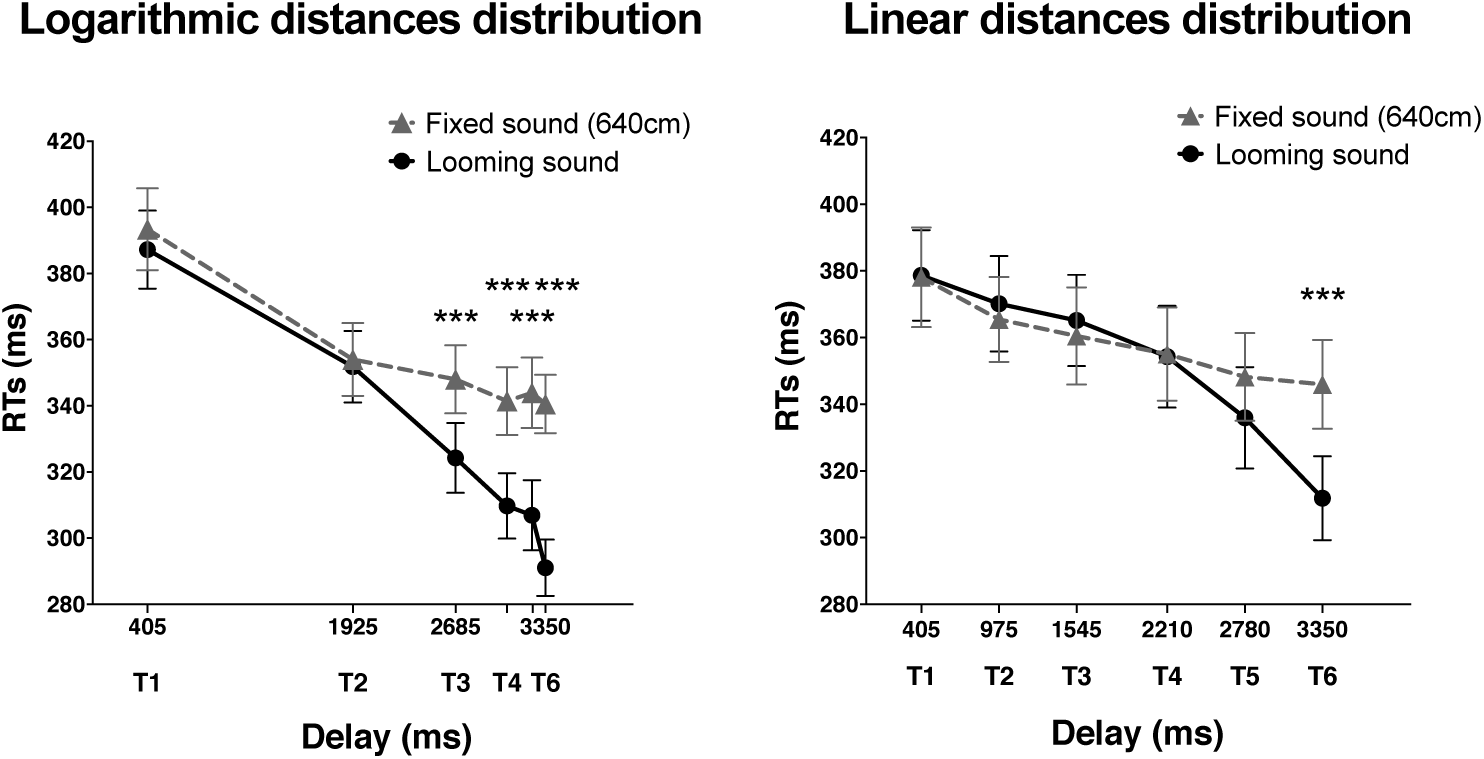
Evaluation of the audiotactile effect on RTs on the linear and logarithmic distances distributions. This figure reports the mean tactile reaction times (±SEM) in presence of the fixed sound (dashed line) or the looming sound (solid line), as a function of the delay of tactile stimulation delivery from the trial beginning. Data of the logarithmic distances distribution session are represented in the left graph, and data for the linear distances distribution session in the right graph. Asterisks indicate significant differences in RTs between fixed sound and looming sound (***p<0.001). On the logarithmic distances distribution, RTs started to be significantly boosted by the proximity of the sound at T3, whereas on the linear distances distribution this significant boost appeared only at T6.

We compared RTs as a function of the presence of the looming or the fixed sound, for each bimodal delay. RTs were significantly faster when the tactile stimulus occurred in presence of a looming sound than in presence of a fixed sound at the delays T3, T4, T5 and T6 (Post-hoc Newman-Keuls’ test: *p* < .001 in all cases). There were no significant difference between RTs in the fixed and looming sound conditions for the delays T1 and T2 (Post-hoc Newman-Keuls’ test: p = .61 and p= .15 respectively). **Linear distances distribution.** We conducted an ANOVA on the mean RTs of the Linear distances distribution only, with the within-subject factors SOUND MOVEMENT (two levels: fixed, looming), and the factor DELAY (six levels: T1, T2, T3, T4, T5 and T6). Analysis revealed a significant main effect of SOUND MOVEMENT (F_(1,17)_ =5.45, p <.05, *η*_*p*_^*2*^ = .243), and a significant main effect of DELAY (F_(5,85)_ = 32.1, p < .001, *η*_*p*_^*2*^ = .654). The analysis also revealed that the two-way interaction SOUND MOVEMENT x DELAY was significant (F_(5,85)_ = 6.81, p < .001, *η*_*p*_^*2*^ = .286), indicating that tactile RTs variation with the temporal delay of tactile delivery from sound onset depended on the movement of the sound (see **Figure 4 right figure**).

We compared RTs in the presence of the looming to RTs in presence of the fixed sound, for each bimodal delays. RTs were significantly faster when the tactile stimulus occurred in presence of a looming sound compared to a fixed sound at the delay T6 (Post-hoc Newman-Keuls’ test: *p* < .001). There were no significant difference between RTs in the fixed and looming sound conditions for the delays T1, T2, T3, T4 and T5 (Post-hoc Newman-Keuls’ test: p >.05 in all cases).

**Description of the spatial dynamic of audiotactile integration behavioral consequences.** We wanted to find the best and simplest description of audiotactile integration behavioral impact on tactile RTs as a function of the distance of the auditory source in space, in order to provide better tools to evaluate and investigate the phenomenon. To isolate the impact of audiotactile integration from expectancy effects on RTs, we subtracted RTs in the fixed sound condition from RTs in the looming sound condition for each participant, for each distances distribution, and for each delay. We hypothesized that RTs evolution would follow a logarithmic law when plotted as a function of the sound source distances from the body. Thus, RTs would vary linearly as a function of the logarithm of the sound source distances. To test this hypothesis, we fitted logarithmic and linear functions into RTs data of the logarithmic and linear distances distribution sessions plotted as a function of the distance of the auditory source at tactile delivery time. We used as the logarithmic function: *y(x) = a.log(x)+ b*, where *x* represented the independent variable (i.e. the distances or logarithm of distances), *y* the dependent variable (i.e. the RTs), *a* the slope and *b* the threshold for x=1. The linear function we used is described by *y(x) = a.x + b* where *a* is the slope and *b* is the intercept at x = 0. As both functions contain two parameters, we can compare the quality of the fitting by directly comparing the root mean square errors values (RMSE).

We fitted RTs data as a function of sound source distances for the logarithmic and linear distances distribution sessions (see **Figure 5, a and b**). For both distances distributions, analysis revealed that the logarithmic function better described participants’ data than linear function (**logarithmic distances distribution**: t(17) = -2.16, p <.05, two-tailed, RMSE_Logaithmic_fitting_=12.7, RMSE_Linear_fitting_ =14.4; **linear distances distribution**: t(17) = -2.24, p <.05, two-tailed, RMSE_Logarithmic_fittingl_= 17.3, RMSE_Linear_fitting_ = 19.3). Thus, the logarithmic function *y(x) = a.log(x) + b* seems to be a better description of RTs evolution than the linear function. Moreover, following these results, data from both conditions can be described by a linear relation between RTs and the logarithm of sound source distances as it is illustrated on **Figure 5, c and d.**

**Figure 5.**
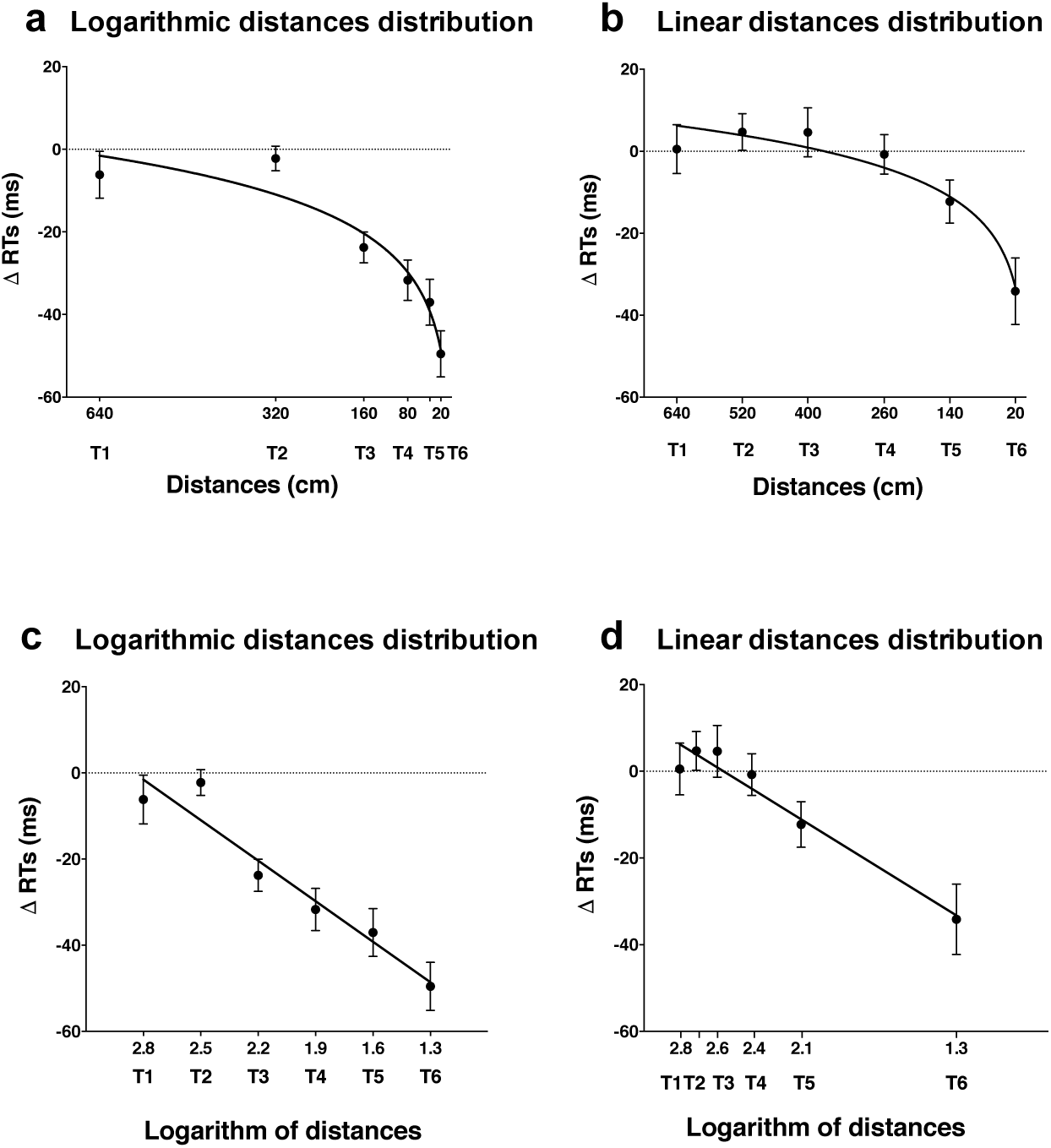
Description of the audiotactile integration consequences on tactile ΔRTs (RTs looming sound – RTs fixed sound) observed in the linear and logarithmic auditory distances distributions sessions. This figure reports the difference of mean tactile reaction times (±SEM) between the two sound movement conditions: the fixed sound and the looming sound. Data of the logarithmic distances distribution condition are represented in the left graphs **(a and c)**, and data for the linear distances distribution condition in the right graphs **(b and d)**. The same data are plotted as a function of the sound source distance (upper graphs, **a and b**) or of the logarithm of the sound source distance (lower graphs, **c and d**). Reaction times are fitted with an increasing logarithmic function when data are plotted as a function of the distance, and with a linear function when data are plotted as a function of the logarithm of the distance.

### 3.2 Auditory Behavioral Assessment Tests

We conducted an ANOVA on participants SUDs, with the within-subject factors DISTANCES DISTRIBUTION (three levels: logarithmic, linear, fixed), and STEP (six levels: S1, S2, S3, S4, S5 and S6). The analysis revealed a significant main effect of DISTANCES DISTRIBUTION (F_(2,38_ =28.9,p< .001, *η _p_* ^*2*^ = .604), of STEP (F_(5,95_ = 67.9, p < .001, *η _p_* ^*2*^ = .781) and a significant interaction between DISTANCES DISTRIBUTION x STEP (F_(10,190)_ = 27.6, p < .001, *η*_*p*_^*2*^ = .592), indicating that SUDs were differently impacted by the steps as a function of the sound source distances distribution.

As illustrated by **Figure 6 panel e**, SUDs did not vary in the **Fixed aBAT**, in which the sound source distance was fixed (Post-hoc Newman-Keuls’ test: p >.70 in all cases). SUDs at 640cm (sound source starting point) and 20cm (sound source ending point) did not significantly differ between the logarithmic and linear aBATs, (Post-hoc Newman-Keuls’ test: p >.40 in both cases). As depicted on **Figure 6, panel a and b,** whereas SUDs significantly increased at each step of the **Logarithmic** a**BAT** (Post-hoc Newman-Keuls’ test: p <.05 in all cases), **in the Linear aBAT,** SUDs did not significantly vary between the first three steps (Post-hoc Newman-Keuls’ test: p > 0.29 in both cases) and started to significantly increase only from S3 (Post-hoc Newman-Keuls’ test: p <.001 in all three cases).

**Figure 6:**
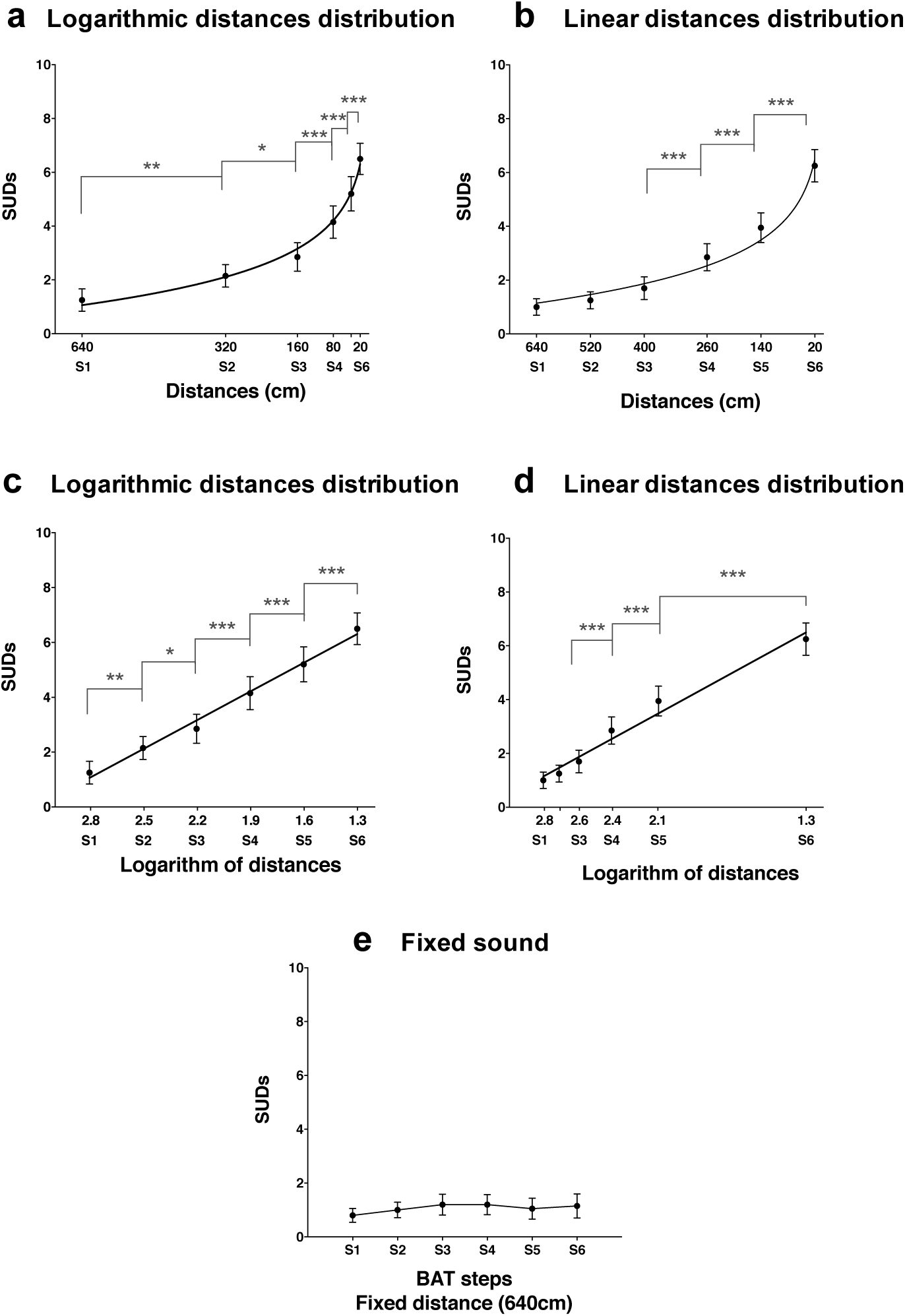
Discomfort rating of aBATS. This figure reports mean SUDs (±SEM) at each step of the three aBATs: the Logarithmic distances aBAT **(a and c)**, the Linear distances aBAT **(b and d)** and the fixed distance aBAT **(e)**. The data of the Logarithmic and Linear aBATs are plotted twice: as a function of the sound source distance (upper graphs, **a and b**) or as a function of the logarithm of the sound source distance (lower graphs, **c and d**). Data of the fixed aBAT are plotted as a function of the steps number, as in this aBAT the sound source distance is fixed at 640cm. Asterisks indicate significant differences in SUDs between distances condition (*p<0.05, **p<0.01, ***p<0.001). SUDs are fitted with a logarithmic function when data are plotted as a function of the distance, and with a linear function when data are plotted as a function of the logarithm of the distance. In the fixed sound condition, there is no evolution of SUDs ratings with the steps (**e**).

Finally, we fitted SUDs data from the logarithmic and linear aBATs as a function of the sound source distances with logarithmic and linear functions. We used as the logarithmic function *y(x) = a.log(x) + b*, where *x* represented the independent variable (i.e. the distances or logarithm of distances), *y* the dependent variable (i.e. the SUDs), *a* the slope and *b* the threshold for *x=*0. The linear function is described by *y(x) = a.x + b* where *a* is the slope and *b* the intercept at *x=0*. As both functions contain two parameters, we can compare the quality of the fitting by directly comparing the root mean square errors values (RMSE). For both conditions, analysis revealed that the logarithmic function described better participants’ data than the linear function (**logarithmic aBAT**: t(19) = -4.9, p <.001, RMSE_Logarithmic_fitting_= 0.58, RMSE_Linear_fitting_ = 1.1; **linear aBAT**: t(19) = -2.3, p <.05, RMSE_Logarithmic_fitting_= 0.55, RMSE_Linear_fitting_ = 0.79). Moreover, following these results, data from both conditions can be described by a linear function between SUDs and the logarithm of sound source distances, as illustrated on **Figure 6, panels c and d.**

## 4. Discussion

We aimed at refining the description of the impact of a sound on tactile detection as a function of the sound source distance, using an audiotactile task based on Canzoneri and colleagues’ paradigm (Canzoneri et al., 2012). We demonstrated that a logarithmic distances distribution was more relevant than a linear one to map events occurring in the auditory space. We also showed that using a condition with a fixed sound located far from participants’ body provided a good baseline to capture expectancy effects. Finally, we fitted the multisensory effects on RTs according to the sound distance to the body and found that a logarithmic decrease was a good descriptor of its variations.

We used as a baseline condition a tactile detection task, in which participants listened to a sound fixed in space, located far from their body. We also studied tactile detection in unimodal conditions, with tactile stimuli occurring before and after the sound (with respectively Tbefore and Tafter). The analysis of unimodal RTs and fixed sound RTs was highly informative. First, RTs decreased with the delay in the fixed sound condition. Furthermore, RTs at T1 and T6 in the fixed sound condition were respectively at the same level than unimodal RTs occurring before and after the sound. Those two observations suggest that, in the fixed sound condition, there is no audiotactile integration effect on RTs and that the observed decrease of RTs with the delay is due to expectation. Subtracting the contribution of expectation to the overall behavioral effect in the looming sound condition gives a properer measure of the behavioral impact of audiotactile integration. In the looming condition, RTs difference between T1 and T6 is about -100ms in the logarithmic session and -67ms in the linear session. Comparing with RTs difference in the fixed sound condition (−49ms in the logarithmic session and -33ms in the linear session), we conclude that at T6, the RTs speeding up accounts for 50% of audiotactile integration effects and for 50% of expectancy effect. This ratio might be dependent on the proportion of catch trials in the experiment (33% here). Furthermore, as predicted, the evolution of expectation is dependent on the distances distribution (Niemi and Naatanen, 1981). Thus, it is important to evaluate expectancy at each delay, in every new experiment.

The majority of previous studies using similar paradigms did not control for expectancy effects (e.g. Canzoneri et al., 2013; Ferri et al., 2015; Taffou and Viaud-Delmon, 2014). We evidenced here that expectation has to been taken into account, as it is responsible for a significant part of the observed behavioral effect. Not assessing the contribution of expectation could be misleading: results attributed to multisensory integration effects could be due to expectancy. Some studies in the literature included a baseline condition: they used unimodal tactile trials, in which tactile stimulations were delivered in the absence of sound. Unimodal tactile trials were usually administered at two different temporal delays, corresponding to the equivalent time of the nearest and the farthest distance used in the audiotactile experiment (for example at T1 and T6 here) (Noel et al., 2015a; Salomon et al., 2017; Serino et al., 2015). Several issues are related to this method. First, expectancy effects may be different in the absence of sound. The sound beginning and end points give participants a temporal window during which the tactile stimulation can occur, thus impacting participants’ anticipation. Second, unimodal trials RTs are indicators of an overall size of expectancy effect but do not evaluate the dynamic changes during the trial. Expectation needs to be evaluated at each delay to reveal the evolution of audiotactile integration impact on RTs according to the sound distance.

Audiotactile integration in the current paradigm depends on the sound dynamic and on the distances distribution. In the logarithmic distances distribution, we observed a speed up effect due to the looming sound presence at the delays T3, T4, T5 and T6. In comparison, the speed up effect appeared only at T6 with the linear distances distribution. This result is not surprising considering that perception of sound source distance is based on a cue varying logarithmically: the largest variation in sound level occurs when the sound is in the nearness of the body. While studies on sound perception in depth are developed using a logarithmic scale of tested distances (Alais and Carlile, 2005; Zahorik and Wightman, 2001), it is remarkable that a paradigm examining the coding of audiotactile stimuli as a function of the distance between the looming sound and the body does not integrate a logarithmic scale of auditory distances. Using a logarithmic distances distribution gives a better resolution of PPS morphometry, using a cognitively relevant mapping of the auditory space.

In the Auditory Behavioral Assessment Tests (aBATs), we collected participants’ emotional experience according to their distance to the sound source and indirectly verified that they perceived the different sound positions. Coherently, SUDs increased as the distance between participants and the sound source decreased, except for the condition in which the sound source did not move. Moreover, similarly to tactile RTs, SUDs evolved logarithmically with the sound source distance. As in the audiotactile test, the aBATs results support the idea that a logarithmic distances distribution is more efficient to evaluate behavioral reactions linked to the location of an auditory stimulation in depth. Further, the logarithmic pattern observed in both tests suggests a pertinent, simpler and parsimonious manner to describe and analyze the data obtained: fitting a linear function on the data plotted as a function of the logarithm of the distance.

To capture the dynamic of multisensory integration according to the distance between the auditory stimulus and the body, we fitted RTs data with linear and logarithmic functions. A linear fitting gave a good description of RTs evolution as a function of the logarithm of the distances. Previous studies proposed a sigmoidal function as a representation of RTs evolution depending on the distance of the sound source. A sigmoidal function is a “S”-shaped curve, in which data evolve between a lower and an upper threshold. The inflexion point, i.e. the central point between both thresholds, is considered as measure of PPS size in space (Canzoneri et al., 2013, 2012). In such a framework, PPS has a precise extent in space, implying that behavioral data follow an in-or-out response. With our data, a sigmoidal fitting was not convincing. Moreover, we wanted to use an economical function in term of parameters (a sigmoidal function has four parameters, the linear and logarithmic functions used here have only two parameters). Our analysis suggested that audiotactile integration behavioral effects follow a logarithmic decrease with the distance (thus a linear decrease with the logarithm of distances). Following this result, multisensory integration behavioral impacts according to the sound distance to the body do not display an in-or-out response, but a gradual response which strength decreases with distance until being null. Thus, PPS might not be an in-or-out zone. In line with this result, previous studies suggested that, within PPS, the distance between an auditory or visual stimuli continues to influence tactile detection (de Haan et al., 2016; Hobeika et al., 2018) and action preparation (Camponogara et al., 2015) This logarithmic function is also coherent with the theoretical framework proposed by Bufacchi and Iannetti, who described PPS as an action field, in which responses are graded with proximity (Bufacchi and Iannetti, 2018).

In this study, we proposed different modifications of the Canzoneri’s audiotactile paradigm to overcome some limitations and improve the power of data analysis. There are still limitations that we did not address. The paradigm is based on the perception of distances. The absolute distance estimation of sound sources is usually a difficult task for non-familiar stimuli in absence of reference, in which participants are not accurate (Middlebrooks and Green, 1991; Zahorik et al., 2005). However, individuals are accurate in the relative comparison of distances between two sources at different distances. The paradigm rests on looming sounds, and compares the effect of continuously varying sound distances. Due to the variability in absolute distances estimation, the method cannot give results in terms of metrical distances but in terms of distances comparisons. Moreover, the results are highly linked to the range of tested distances (Poulton, 1975). Subjects learn the range of stimuli used in the experiment and adapt their behaviors to it. It would be deceptive to give metrical estimation of PPS considering the influence of range effects.

Even if we focused in our work on mastering the auditory aspects of the audiotactile paradigm, the importance of the selection of tactile stimulation type requires to be emphasized. Human skin tactile properties vary widely between different body parts (Chouvardas et al., 2008; Dargahi and Najarian, 2004). In the present study, we decided to stimulate finger pads because it is one of the most sensitive body part, with a high density of tactile receptors and a good spatial resolution (Johansson and Vallbo, 1979a, 1979b). Our tactile device was a mechanical tactile device: a miniature speaker able to send fast stimulations (20ms in the experiment), and which can send all forms of signal. Electric stimulation can also be used. Electric shocks are detected easily with a good spatial resolution, but they can be painful and non-ecological (Chouvardas et al., 2008). Further refining the audiotactile paradigm implicates identifying the type of tactile stimulation that is the most suited to the experimental context, and to the targeted body part and its tactile receptors.

## 5. Conclusion

It is important to master every aspect of the auditory and tactile stimulations to develop a reliable and efficient protocol to study peripersonal space. For auditory stimulations in depth, distances distribution needs to be logarithmic to have a relevant mapping of the space. After controlling for expectancy effects, we found that audiotactile behavioral impacts change logarithmically with sound distance. This finding has important implications for the theoretical aspects of PPS: behavioral responses linked to PPS coding do not follow an in-or-out pattern but a rather gradual pattern. Furthermore, this efficient method to describe audiotactile integration in space could lead to more reliable and precise results for studies aiming at determining the interplay between PPS and multisensory integration.

## Acknowledgments

We are grateful to Emmanuel Fléty and Arnaud Recher for their help with the apparatus for tactile stimulation. This work was supported by Sorbonne Universités Investissements d’avenir, Emergence and the French Ministry of Armed Forces (grant number HUM-1-0820).

## Footnotes

1 The linear distances distribution is actually pseudo-linear. The spaces between tested distances are not strictly regular: the space between T1-T2, T2-T3, T4-T5, and T5-T6 is 120cm., whereas the space between T3-T4 is 140cm. This difference is due to experimental design constraints

## Bibliography

Alais, D., Carlile, S., 2005. Synchronizing to real events: subjective audiovisual alignment scales with perceived auditory depth and speed of sound. Proc. Natl. Acad. Sci. U. S. A. 102, 2244–7. https://doi.org/10.1073/pnas.0407034102

Bernasconi, F., Noel, J.P., Park, H.D., Faivre, N., Seeck, M., Spinelli, L., Schaller, K., Blanke, O., Serino, A., 2018. Audio-tactile and peripersonal space processing around the trunk in human parietal and temporal cortex: An intracranial EEG study. Cereb. Cortex 28, 3385–3397. https://doi.org/10.1093/cercor/bhy156

Bufacchi, R.J., Iannetti, G.D., 2018. An Action Field Theory of Peripersonal Space. Trends Cogn. Sci. xx, 1–15. https://doi.org/10.1016/j.tics.2018.09.004

Camponogara, I., Komeilipoor, N., Cesari, P., 2015. When distance matters: Perceptual bias and behavioral response for approaching sounds in peripersonal and extrapersonal space. Neuroscience 304, 101–108. https://doi.org/10.1016/j.neuroscience.2015.07.054

Canzoneri, E., Magosso, E., Serino, A., 2012. Dynamic sounds capture the boundaries of peripersonal space representation in humans. PLoS One 7, e44306. https://doi.org/10.1371/journal.pone.0044306

Canzoneri, E., Ubaldi, S., Rastelli, V., Finisguerra, A., Bassolino, M., Serino, A., 2013. Tool-use reshapes the boundaries of body and peripersonal space representations. Exp. Brain Res. 228, 25–42. https://doi.org/10.1007/s00221-013-3532-2

Cardini, F., Fatemi-ghomi, N., Gajewska-knapik, K., Gooch, V., Aspell, J.E., 2019. Enlarged representation of peripersonal space in pregnancy. Sci. Rep. 9, 1–7. https://doi.org/10.1038/s41598-019-45224-w

Carpentier, T., Noisternig, M., Warusfel, O., 2015. Twenty years of Ircam Spat: looking back, looking forward. Int. Comput. Music Conf. Proc. 270–277.

Chouvardas, V.G., Miliou, A.N., Hatalis, M.K., 2008. Tactile displays: Overview and recent advances. Displays 29, 185–194. https://doi.org/10.1016/j.displa.2007.07.003

Cléry, J., Ben Hamed, S., 2018. Frontier of self and impact prediction. Front. Psychol. 9, 1–17. https://doi.org/10.3389/fpsyg.2018.01073

Cléry, J., Guipponi, O., Odouard, S., Wardak, C., Ben Hamed, S., Clery, J., Guipponi, O., Odouard, S., Wardak, C., Ben Hamed, S., 2015. Impact Prediction by Looming Visual Stimuli Enhances Tactile Detection. J. Neurosci. 35, 4179–4189. https://doi.org/10.1523/JNEUROSCI.3031-14.2015

Colby, C.L., Duhamel, J.-R., Goldberg, M.E., 1993. Ventral intraparietal area of the macaque: anatomic location and visual response properties. J. Neurophysiol. 69, 902–14.

Dargahi, J., Najarian, S., 2004. Human tactile perception as a standard for artificial tactile sensing — a review. Int. J. Med. Robot. Comput. Assist. Surg. 1, 23–35. https://doi.org/10.1581/mrcas.2004.010109

de Haan, A.M., Smit, M., Van der Stigchel, S., Dijkerman, H.C., 2016. Approaching threat modulates visuotactile interactions in peripersonal space. Exp. Brain Res. 234, 1875–1884. https://doi.org/10.1007/s00221-016-4571-2

Farnè, A., Làdavas, E., 2002. Auditory peripersonal space in humans. J. Cogn. Neurosci. 14, 1030–43. https://doi.org/10.1162/089892902320474481

Ferri, F., Tajadura-Jiménez, A., Väljamäe, A., Vastano, R., Costantini, M., 2015. Emotion-inducing approaching sounds shape the boundaries of multisensory peripersonal space. Neuropsychologia 70, 468–475. https://doi.org/10.1016/j.neuropsychologia.2015.03.001

Fontana, F., Rocchesso, D., 2008. Auditory distance perception in an acoustic pipe. ACM Trans. Appl. Percept. 5, 1–15. https://doi.org/10.1145/1402236.1402240

Gentilucci, M., Fogassi, L., Luppino, G., Matelli, M., Camarda, R., Rizzolatti, G., 1988. Functional organization of inferior area 6 in the macaque monkey - I. Somatotopy and the control of proximal movements. Exp. Brain Res. 71, 475–490.

Graziano, M.S.A., Cooke, D.F., 2006. Parieto-frontal interactions, personal space, and defensive behavior. Neuropsychologia 44, 2621–2635. https://doi.org/10.1016/j.neuropsychologia.2005.09.011

Graziano, M.S.A., Gross, C.G., 1993. A bimodal map of space: somatosensory receptive fields in the macaque putamen with corresponding visual receptive fields. Exp. Brain Res. 97, 96–109.

Hershenson, M., 1962. Reaction time as a measure of intersensory facilitation. J. Psychol. 63, 289–293. https://doi.org/10.1037/h0055703

Hobeika, L., Taffou, M., Viaud-Delmon, I., 2019. Social coding of the multisensory space around us. R. Soc. Open Sci. 6, 181878.

Hobeika, L., Viaud-Delmon, I., Taffou, M., 2018. Anisotropy of lateral peripersonal space is linked to handedness. Exp. brain Res. 236, 609–618. https://doi.org/10.1007/s00221-017-5158-2

Johansson, R.S., Vallbo, A.B., 1979a. Tactile sensibility in the human hand: Relative and absolute densities of four types of mechanoreceptive units in glabrous skin. J. Physiol. 286, 283–300.

Johansson, R.S., Vallbo, A.B., 1979b. Detection of tactile stimuli. Thresholds of afferent units related to psychophysical thresholds in the human hand. J. Physiol. 297, 405–422. https://doi.org/10.1113/jphysiol.1979.sp013048

Kandula, M., Hofman, D., Dijkerman, H.C., 2015. Visuo-tactile interactions are dependent on the predictive value of the visual stimulus. Neuropsychologia 70, 358–366. https://doi.org/10.1016/j.neuropsychologia.2014.12.008

Kandula, M., Van der Stoep, N., Hofman, D., Dijkerman, H.C., 2017. On the contribution of overt tactile expectations to visuo-tactile interactions within the peripersonal space. Exp. Brain Res. 235, 2511–2522. https://doi.org/10.1007/s00221-017-4965-9

Kolarik, A.J., Moore, B.C.J., Zahorik, P., Cirstea, S., 2015. Auditory distance perception in humans: a review of cues, development, neuronal bases, and effects of sensory loss. Attention, Perception, Psychophys. 78, 373–395. https://doi.org/10.3758/s13414-015-1015-1

Làdavas, E., Farnè, A., 2004. Visuo-tactile representation of near-the-body space. J. Physiol. Paris 98, 161–170.

Ladavas, E., Pavani, F., Farnè, A., 2001. Auditory Peripersonal Space in Humans: a Case of Auditory-Tactile Extinction. Neurocase 7, 97–103. https://doi.org/10.1093/neucas/7.2.97

Lang, P.J., Lazovik, A.D., 1963. Experimental desensitization of a phobia. J. Abnorm. Soc. Psychol. 66, 519–25.

Maravita, A., Spence, C., Driver, J., 2003. Multisensory integration and the body schema: Close to hand and within reach. Curr. Biol. 13, 531–539. https://doi.org/10.1016/S0960-9822(03)00449-4

Middlebrooks, J.C., Green, D.M., 1991. Sound Localization by Human Listeners. Annu. Rev. Psychol. 42, 135–159. https://doi.org/10.1146/annurev.ps.42.020191.001031

Nicholls, M.E.R., Thomas, N.A., Loetscher, T., Grimshaw, G.M., 2013. The flinders handedness survey (FLANDERS): A brief measure of skilled hand preference. Cortex 49, 2914–2926. https://doi.org/10.1016/j.cortex.2013.02.002

Niemi, P., Naatanen, R., 1981. Foreperiod and Simple Reaction Time. Psychol. Bull. 89, 133–162. https://doi.org/10.1037/0033-2909.89.1.133

Noel, J.-P., Grivaz, P., Marmaroli, P., Lissek, H., Blanke, O., Serino, A., 2015a. Full body action remapping of peripersonal space: The case of walking. Neuropsychologia 70, 375–384. https://doi.org/10.1016/j.neuropsychologia.2014.08.030

Noel, J.-P., Pfeiffer, C., Blanke, O., Serino, A., 2015b. Peripersonal space as the space of the bodily self. Cognition 144, 49–57. https://doi.org/10.1016/j.cognition.2015.07.012

Poulton, E.C., 1975. Range Effects in Experiments on People. Am. J. Psychol. 88, 3–32.

Rizzolatti, G., Scandolara, C., Matelli, M., Gentilucci, M., 1981. Afferent properties of periarcuate neurons in macaque monkeys. I. Somatosensory responses. Behav. Brain Res. 2, 125–146. https://doi.org/10.1016/0166-4328(81)90052-8

Salomon, R., Noel, J.P., Lukowska, M., Faivre, N., Metzinger, T., Serino, A., Blanke, O., 2017. Unconscious integration of multisensory bodily inputs in the peripersonal space shapes bodily self-consciousness. Cognition 166, 174–183. https://doi.org/10.1016/j.cognition.2017.05.028

Serino, A., Noel, J.-P., Galli, G., Canzoneri, E., Marmaroli, P., Lissek, H., Blanke, O., 2015. Body part-centered and full body-centered peripersonal space representation. Sci. Rep. 5, doi:10.1038/srep18603. https://doi.org/10.1038/srep18603

Shinn-Cunningham, B.G., 2000. Distance Cues for Virtual Auditory Space. Proc. IEEE-PCM 2000, 227–230. https://doi.org/10.1.1.73.2560

Spence, C., Nicholls, M.E., Gillespie, N., Driver, J., 1998. Cross-modal links in exogenous covert spatial orienting between touch, audition, and vision. Percept. Psychophys. 60, 544–557. https://doi.org/10.3758/BF03206045

Sumby, W.H., Pollack, I., 1954. Visual Contribution to Speech Intelligibility in Noise. J. Acoust. Soc. Am. 26, 212–215. https://doi.org/10.1121/1.1907309

Taffou, M., Viaud-Delmon, I., 2014. Cynophobic Fear Adaptively Extends Peri-Personal Space. Front. Psychiatry 5, 122. https://doi.org/10.3389/fpsyt.2014.00122

Tajadura-Jiménez, A., Väljamäe, A., Asutay, E., Västfjäll, D., 2010. Embodied auditory perception: The emotional impact of approaching and receding sound sources. Emotion 10, 216–229. https://doi.org/10.1037/a0018422

Thyer, B.A., Papsdorf, J.D., Davis, R., Vallecorsa, S., 1984. Autonomic correlates of the subjective anxiety scale. J. Behav. Ther. Exp. Psychiatry 15, 3–7.

Van Bockstaele, B., Verschuere, B., Koster, E.H.W., Tibboel, H., De Houwer, J., Crombez, G., 2011. Effects of attention training on self-reported, implicit, physiological and behavioural measures of spider fear. J. Behav. Ther. Exp. Psychiatry 42, 211–218. https://doi.org/10.1016/j.jbtep.2010.12.004

Van der Biest, L., Legrain, V., De Paepe, A., Crombez, G., 2016. Watching what’s coming near increases tactile sensitivity: An experimental investigation. Behav. Brain Res. 297, 307–314. https://doi.org/10.1016/j.bbr.2015.10.028

Wolpe, J., 1973. The practice of behavior therapy. Pergamon.

Zahorik, P., Brungart, D.S., Bronkhorst, A.W., 2005. Auditory Distance Perception in Humans: A Summary of Past and Present Research. Acta Acust. united with Acust. 91, 409–420.

Zahorik, P., Wightman, F.L., 2001. Loudness constancy with varying sound source distance. Nat. Neurosci. 4, 78–83. https://doi.org/10.1038/82931

